# Intermediate abundance promotes speciation when dispersal is limited

**DOI:** 10.64898/2026.05.22.727295

**Authors:** Andrew J. Rominger, Daniel S. Gruner, Isaac Overcast, James L. Rosindell, Catherine E. Wagner

## Abstract

Why do some lineages diversify while others do not? This remains a central question in evolutionary ecology. A long-standing assumption, dating to Darwin and embedded in the Unified Neutral Theory of Biodiversity, holds that abundant species should speciate at higher rates. Conversely, theoretical and empirical work highlights the possibility that rare and dispersal-limited clades might be more prone to speciation. Using a birth-death-immigration model with protracted speciation in a multi-population landscape connected by limited dispersal, we show that abundance has a hump-shaped effect on probability of speciation. Our model reveals that intermediate abundance maximizes speciation probability because larger populations disperse more, swamping regional differentiation and inhibiting speciation completion, while smaller populations lack the persistence and incipient speciation needed to diversify. We find empirical support for this prediction with an analysis of data from arthropods endemic to Hawai’i, where genus-level species richness shows a significant hump-shaped relationship with mean genus abundance. These findings provide a mechanistic explanation for a nuanced relationship between abundance and diversification.

## Introduction

A fundamental question in evolutionary ecology concerns why some clades are diverse while others are not. Are there properties of species and lineages that promote speciation? One line of inquiry with a long and convoluted history is the role of commonness and rarity in the process of diversification. Going back over 150 years, Darwin (1859) argued that widespread, abundant species should be superior competitors and thus more likely to give rise to new species.

Building on this foundation, Yule’s (1925) model, which became a landmark contribution to modeling diversification, was initially intended as a way to incorporate demography into predictions for diversification and clade richness (Pennell and MacPherson 2025). Despite these deep roots in evolutionary thinking, little consensus has emerged on the relationship between abundance and diversification. Darwin’s line of thinking carried forward to macroecologists who attempted to model, with some empirical support, the presumed positive association between commonness, fitness, and speciation (Maurer 1999).

In parallel, paleontologists searched for relationships between commonness and diversification, alternately finding evidence for a positive relationship (Krug et al. 2008) but also a negative relationship (Stanley 1986, Jablonski and Roy 2003). Empirical studies of modern taxa have largely found that commonness is negatively or inconclusively related to measures of diversification (Jablonski and Roy 2003, Makarieva and Gorshkov 2004, Greenberg and Mooers 2017, Smyčka et al. 2023, Afonso Silva et al. 2025, but see Hay et al. 2022). While some of these studies focus specifically on average abundance, others on range size, and some on both, the two macroecological properties are most often strongly correlated, a fascinating macroecology pattern in and of itself (Brown 1995, Gaston 2003). This correlation is no doubt caused, at least in part, by the net increase in dispersal of larger populations owing simply to chance: if each individual carries a probability of dispersing, more individuals in a population will lead to more overall dispersal (Hubbell 2001). It is also of course possible that shared underlying mechanisms cause both increased population growth rate and increased dispersal rate (Gaston et al. 1997, Holt et al. 1997).

More recent theoretical and modeling work has largely assumed a positive relationship between commonness and diversification. Hubbell (2001) perhaps started this trend with the inclusion of “point mutation” speciation in the Unified Neutral Theory of Biodiversity (UNTB). Because point mutation speciation assumes a constant probability of speciation for every individual, lineages with more individuals will experience more speciation events in the UNTB (Hubbell 2001, Etienne et al. 2007). The speciation mechanism first proposed by Hubbell (2001) was never an accurate model of real diversification (Ricklefs 2003), but even later attempts to increase the realism of speciation in the UNTB retain the emergent property that more abundant lineages will undergo more speciation. Protracted speciation (Rosindell et al. 2010) again assumes that incipient speciation is constant across individuals and thus full speciation will scale positively with lineage abundance; fission speciation (Etienne and Haegeman 2011) also assumes the probability of fission (the event that leads to speciation) increases with the number of individuals in a species. Largely independently from UNTB and its descendants, phylogenetic models of geographic change and diversification also explicitly assume that more widespread species have greater opportunity for speciation (Goldberg et al. 2011).

The process that underlies speciation itself may suggest something different in terms of the relationship between speciation, commonness, and rarity: dispersal. Dispersal is a key mechanism by which populations can become isolated or connected, potentially leading to speciation or admixture (Yamaguchi 2022). Dispersal figures prominently in phylogenetic models of the geographic range and diversification (Matzke 2014), it is necessary to maintain local biodiversity in the UNTB (Hubbell 2001), and it is the mechanism connecting abundance to range size in both neutral (Hubbell 2001) and non-neutral (Brown 1995) macroecological models.

Due to the potential for dispersal to both create population isolates but also swamp out regional differences that could have led to speciation, the connection between dispersal ability as a biological trait and speciation as an evolutionary outcome remains a fertile line of inquiry. Some empirical studies have found that measures of diversification have a hump-shaped or negative relationship with morphological proxies for dispersal ability (Price and Wagner 2004, Claramunt et al. 2012, Czekanski-Moir and Rundell 2019) while others have found a positive to flat relationship between dispersal ability and diversification (Claramunt et al. 2025). Agent-based model simulations have supported the idea of a trade-off between too little dispersal leading to population instability and too much dispersal leading to admixture with intermediate dispersal balancing the two extremes allowing for speciation (Birand et al. 2012, Ashby et al. 2020, Ciccheto et al. 2024).

But dispersal is not just a trait connected to morphology or imposed in a simulation, it emerges from population dynamics: each individual carries some probability of dispersing, thus species with more individuals will present with higher dispersal. This is the assumption, a realistic one, in birth-death-immigration models (Kendall 1948). Therefore, commonness and dispersal are interconnected and interdependent in their effects on speciation if they indeed have any consistent effects. It also remains an open debate whether their effects result from the determinism of adaptive evolution as Darwin (1859) believed or instead emerge from chance.

Here we develop a birth-death-immigration model with protracted speciation embedded in a landscape of multiple local populations connected with limited dispersal to investigate the role of abundance and stochastic dispersal in modulating the probability of speciation. Contrary to Darwin (1859) and UNTB (Hubbell 2001, Etienne et al. 2007, Rosindell et al. 2010, Etienne and Haegeman 2011), we find that intermediate abundances lead to the greatest probability for speciation. Critically, this result does not depend in any way on whether populations are more or less adaptively fit, it only depends on a balance between large enough population sizes to accrue sufficient probability of incipient speciation but small enough population sizes to not lose regional differentiation due to increased dispersal. We also analyze real data on the species richness and abundance of arthropod communities in Hawai’i finding empirical evidence for intermediate abundance promoting speciation. The isolation of Hawai’i, leading to tractable endemic radiations, and its known geologic age, ranging from still forming to 4–6 million years, make it an ideal study system for investigating the interplay of abundance and speciation.

## Methods

### Simulating a birth-death-immigration process with speciation

We simulate a birth-death-immigration process with speciation in a modified metapopulation (Hanski 1998) setting. For simplicity, we consider only a single species (not multi-species) metapopulation. In this setting there are a number of local populations (minimum two are needed, we simulate 10) fully connected by local dispersal as well as a global source population connected to each local population by dispersal. The global source population is considered extremely large relative to local populations such that extinction of the global source population is unlikely on the time scale of local dynamics (we approximate this assumption by never allowing the source population to go extinct). Local populations start at size 0 and grow and shrink according to local births and deaths, dispersal between local populations, and dispersal from the source pool. A new species arise via a modified protracted speciation process (Rosindell et al. 2010) similar to the model of Tao et al. (2021). Our simulation model is consistent with other models derived from the UNTB (Hubbell 2001, Rosindell and Harmon 2013, Rosindell et al. 2015), but unlike the multi-species perspective of those models, we concern ourselves only with a single species and whether that one species undergoes a speciation event. In this way we do not make the assumption of per capita neutrality across species, meaning our results could generalize to multi-species eco-evolutionary models based on niche theories [e.g. Tilman (1982); Chesson (2000); letten2017].

Our simulation proceeds according to these rules:

1. Local populations are initialized as empty.
2. Per capita birth, death, immigration, and speciation all happen independently and are determined by their own respective rates.
3. Speciation has 2 steps:
  i. Incipient speciation occurs at the individual level and its population is flagged as containing an incipient species. Subsequent incipient speciation events in the same local population are ignored.
  ii. If the incipient species survives long enough it becomes a new species; “long enough” is determined by a waiting time parameter τ—the larger τ, the longer the time frame for a fully new species to emerge.
4. Dispersal between local communities and from the global source pool slows the progress toward speciation; if an incipient species has to wait τ time (in the absence of immigration) until it is a full species, each immigration event adds an increment to τ of *ξ*/*n*_*i*_ where *ξ* is a parameter we can set and *n*_*i*_ is the size of the population containing the incipient species. If an individual of an incipient species disperses to a different population, it is subsumed within that population and its incipient status is lost.
5. If a local population goes extinct, the speciation tracking for that patch is reset.
6. The simulation is stopped under one of two conditions: 1) if any incipient species becomes a fully diverged new species, or 2) if the simulation reaches the maximum designated number of iterations.

Because of the stopping conditions, each simulation replicate will either yield one and only one full speciation, or no full speciation events at all. We can thus estimate the probability of speciation as the proportion of simulations that yield a full speciation event.

Further details of the simulation are provided in Supplementary Section S2. All simulation code is written in R (R Core Team 2025) using *Rcpp* (Eddelbuettel et al. 2025a) and *RcppArmadillo* (Eddelbuettel et al. 2025b). Underlying C++ and R source code is available in the R package *abundolism* (Rominger 2026) accompanying this paper.

### Simulation experiment

Table 1 summarizes the simulation parameters in our model and the range of values we use in our simulation experiment. In all cases parameter values are drawn from a uniform distribution. A total of 5,000 simulations were run, each with a unique set of randomly sampled parameters. Parameter values and the number of iterations were chosen with three considerations in mind:

**Table 1:** Parameters governing the simulation and their ranges over the simulation experiment. Mathematical parameter names are given first followed by their name in the simulation code. Note np and nstep were fixed throughout the course of the simulation experiment.

| parameter |  | range |  | description |
| --- | --- | --- | --- | --- |
| $\lambda$ | (**la**) | 0.1 | 4.0 | per capita local birth rate |
| $\mu$ | (**mu**) | 0.1 | 4.0 | per capita local death rate |
| $\gamma$ | (**g**) | 0.1 | 1.0 | dispersal rate from the global source pool to one local population |
| $m_{\text{prop}}$ | (**m_prop**) | 0.0 | 0.1 | proportional dispersal rate between local populations relative to global rate |
| $\nu$ | (**nu**) | 0.0 | 0.1 | per capita incipient speciation rate |
| $\tau$ | (**tau**) | 20.0 | 200.0 | wait time to full speciation in the absence of dispersal |
| $\xi$ | (**xi**) | 10.0 | 100.0 | amount each dispersal event sets back the progression toward speciation |
| $n_p$ | (**np**) | 10.0 | 10.0 | number of local populations |
| $n_{\text{step}}$ | (**nstep**) | 10,000.0 | 10,000.0 | number of simulation iterations |

1. speciation should be possible to observe in a reasonable number of simulation iterations for computational practicality
2. population sizes should be within realistic ranges
3. timescales should be within reasonable bounds

In the context of the speciation-related parameter values we used, timescales should be interpreted in units of 10,000 years. Thus the maximum value of τ is 2,000,000 years and the maximum rate of incipient speciation *v* is 0.1 per individual per 10,000 years, leading to rates of new species divergence roughly in line with published estimates (Hedges et al. 2015).

Parameter values were chosen as a balance between biological realism and computational efficiency: rates are sufficiently fast to allow for abundances to reach levels often found in survey data (e.g. the arthropod survey data we analyze here) within 10,000 time steps.

### Analysis of arthropod richness and abundance from Hawai’i

To bring real data to bear on our insights from simulations, we analyze the species richness and mean species abundance of arthropod genera in Hawai’i. Although only an approximation, most arthropod genera only arrived in Hawai’i once, meaning all endemic species of those genera represent *in situ* diversification (Cowie and Holland 2008). For the purpose of our analysis, we proceed with this approximation.

To calculate species richness per genus we use the Bernice Pauahi Bishop Museum’s checklist of arthropods in Hawai’i (Nishida 1992). To estimate mean abundance per genus we use an extensive survey of arthropods from across the Hawaiian Archipelago (Gruner 2007). The survey data from Gruner (2007) comprise sample locations from 45 trees across 5 sites. Genus abundance was estimated as the mean abundance of all species belonging to each genus across all sample locations.

We are interested in determining if the number of species per genus (a proxy for diversification) peaks at intermediate genus abundances. To do this we fit a quadratic generalized linear model with species richness as response and abundance as explanatory variable. But the question remains, what distribution should we assume for number of species? Kendall (1948) showed that a birth-death process will result in a geometric distribution of descendants. Conditional on lineage survival at time *t*, we can parameterize the geometric probability mass function as

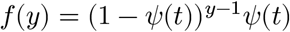

where *y* is the number of species in a lineage. 1 − *ψ*(*t*) is the joint probability of two events: 1) a bifurcation occurs prior to time *t*, and 2) descendants of that bifurcation event being survive to time *t* (Nee et al. 1994). We will call 1 − *ψ*(*t*) the “observable diversification probability.” *ψ*(*t*) correspondingly refers to the probability of survival without further diversification. *ψ*(*t*) depends on rates of speciation and extinction in addition to time *t*.

The number of species in any given lineage *i* extant at time *t* will follow a geometric distribution with probability parameter *p*_*i*_ = *ψ*_*i*_(*t*_*i*_). But we do not know *p*_*i*_ because we do not know the arrival times of most arthropod lineages in Hawai’i—thus do not know *t*_*i*_—and we do not know their rates of speciation and extinction. However, we can model our uncertainty in *p*_*i*_ by letting it be a random variable drawn from a beta distribution. This results in the beta-geometric distribution with parameters *α* and *β*. Interestingly enough, the Yule-Simons distribution, historically developed to model number of species in genera (Yule 1925), is a special case of the beta-geometric distribution when *β* = 1 (Irwin 1975), so our approach has precedent. We will allow *β* to be a free parameter, thus not making the strict birth-only assumption of Yule (1925)

Our final, quadratic model relating genus richness to genus abundance is

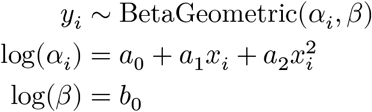

where *x*_*i*_ is average abundance of lineage *i*. This model means that we allow the *α* parameter of the beta geometric distribution to change with lineage abundance but we assume the *β* parameter is fixed across lineages. A fixed *β* across lineages still allows sufficient flexibility of the model without introducing additional parameters for which there may be insufficient power to reliably estimate. From this model our estimate for the observable diversification probability for lineage *i* is 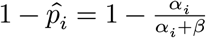.

To evaluate the significance of an intermediate peak in diversification across abundances (i.e. the quadratic term) we also fit a model without quadratic terms:

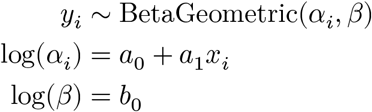

We then compare quadratic and linear models with a likelihood ratio test to evaluate significance of the intermediate peak.

While the *VGAM* package (Yee 2025) in R enables beta-geometric generalized linear modeling, its implementation is based on iterative Fisher scoring which is unnecessary and computationally slow for our application. Therefore, we use functionality from *VGAM* to write a custom likelihood maximization routine (Rominger 2026) that we use to estimate model parameters. More details on this approach are provided in Supplementary Section S5.

## Results

### Simulating a birth-death-immigration process with speciation

Supplementary Section S4 provides details on the outcomes of the simulation experiment, in particular documenting the effect of differing values for speciation rate, dispersal rate, and waiting time to full speciation

### Simulation experiment: effect of abundance on speciation

In visualizing simulation runs that did and did not produce speciation events, we can see a clear peak in the probability of speciation at intermediate abundance Figure 2. A logistic regression model with a quadratic term confirms this peak, finding that the coefficient on the quadratic term falls within the 95% confidence interval (-0.416, -0.365)

**Figure 1:**
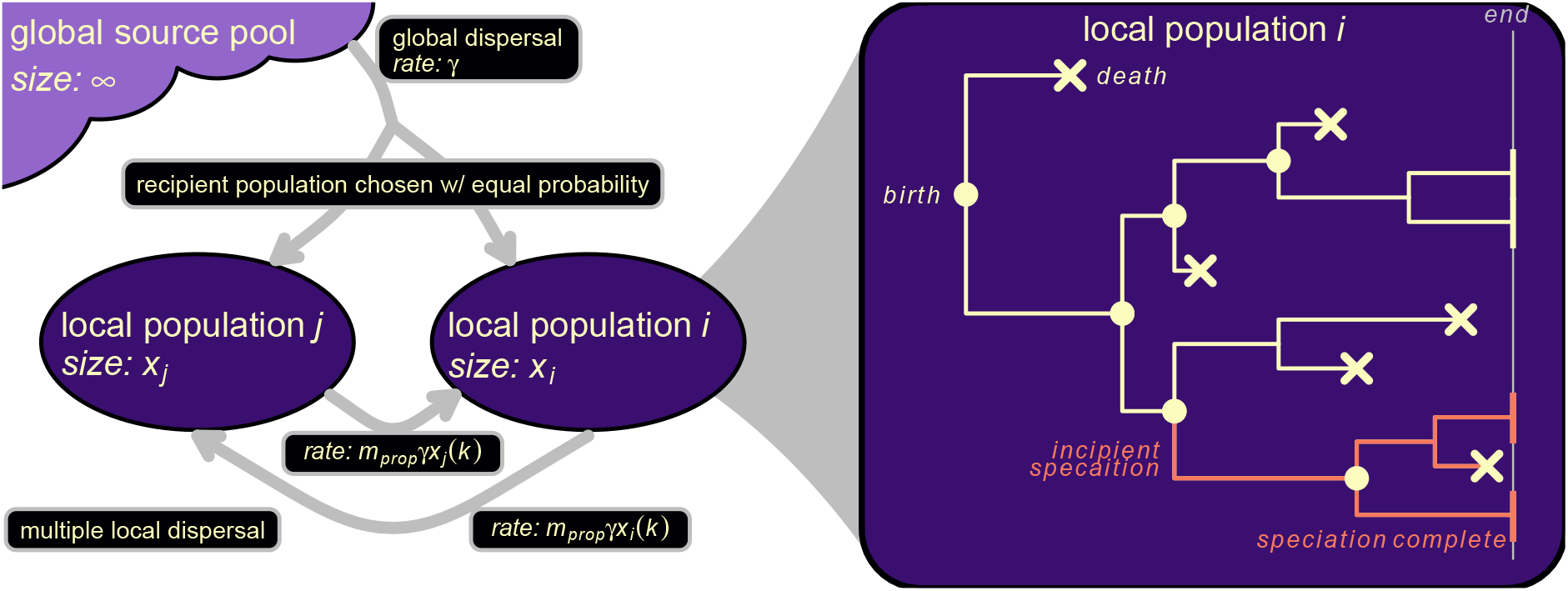
Conceptual overview of our simulation model. Local populations are connected by immigration between populations and also from a global source pool. Within local populations birth and death determine population growth and decline, augmented by immigration. Random incipient speciation events can occur but do not lead to full speciation unless the lineage persists for at least a duration of τ (possibly more in the face of immigration). If full speciation occurs the simulation stops. If no speciation occurs but the maximum number of iterations is reached, the simulation likewise stops.

**Figure 2:**
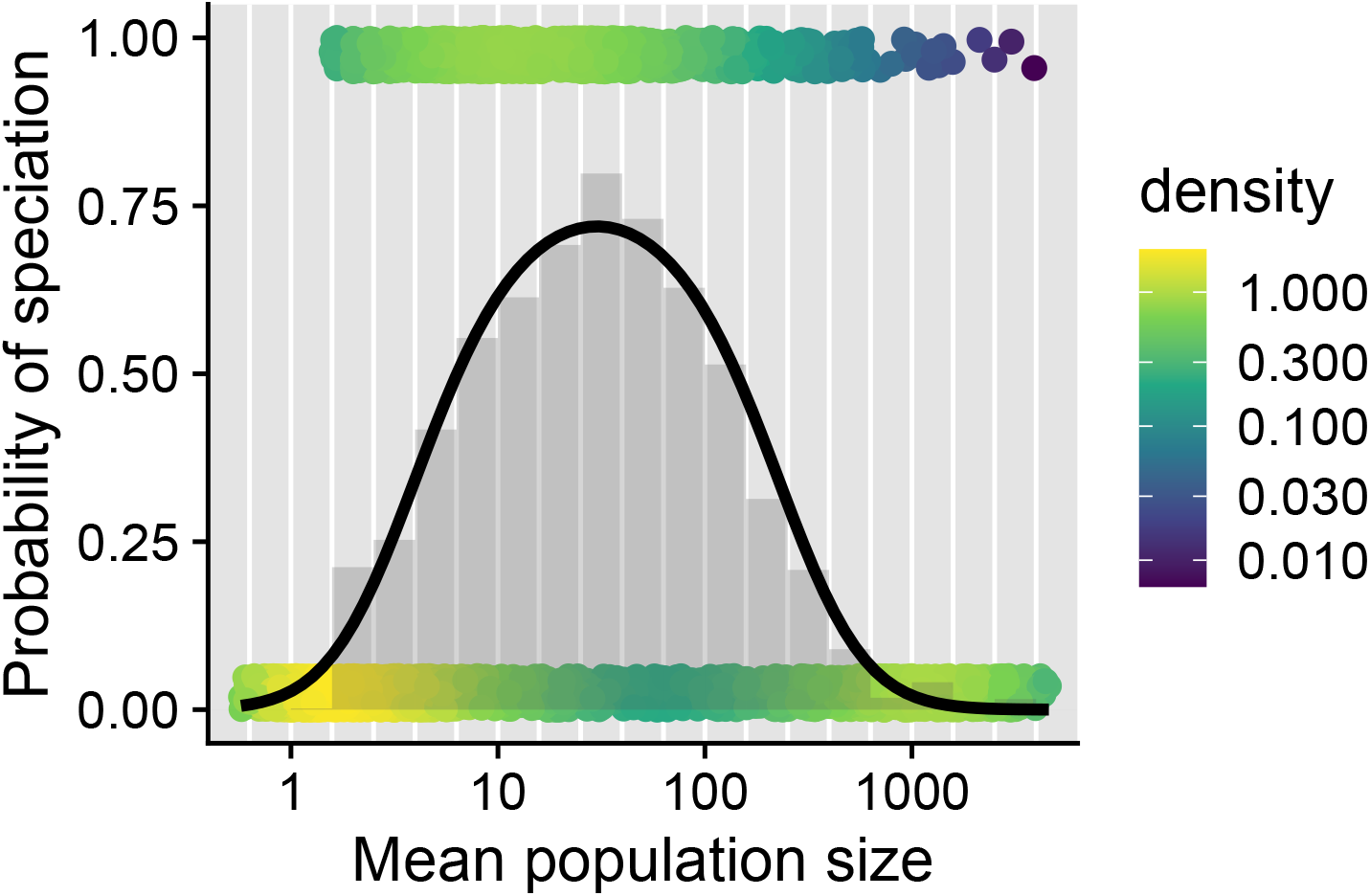
Relationship across simulation runs between average abundance and speciation. Points individual simulation outcomes and are jittered around 1 (yes speciation) and 0 (no speciation). Points are color coded according to local density of points to better show the distribution of simulation results. Gray bars are proportion of simulations that produced speciation binned by mean population size. Curve is a quadratic binomial generalized linear model.

### Relationship between genus abundance and species richness in Hawai’i arthropods

Arthropods in Hawai’i show a hump-shaped relationship between abundance and species richness Figure 3. There is a clear outlier with higher genus species richness (above 300) than expected based on abundance. This outlier is *Drosophila*, which is not surprising given both the extensive systematics work on Hawaiian *Drosophila* and their unique speciation mechanisms (O’Grady and DeSalle 2018), both of which could inflate the group’s reported species richness.

**Figure 3:**
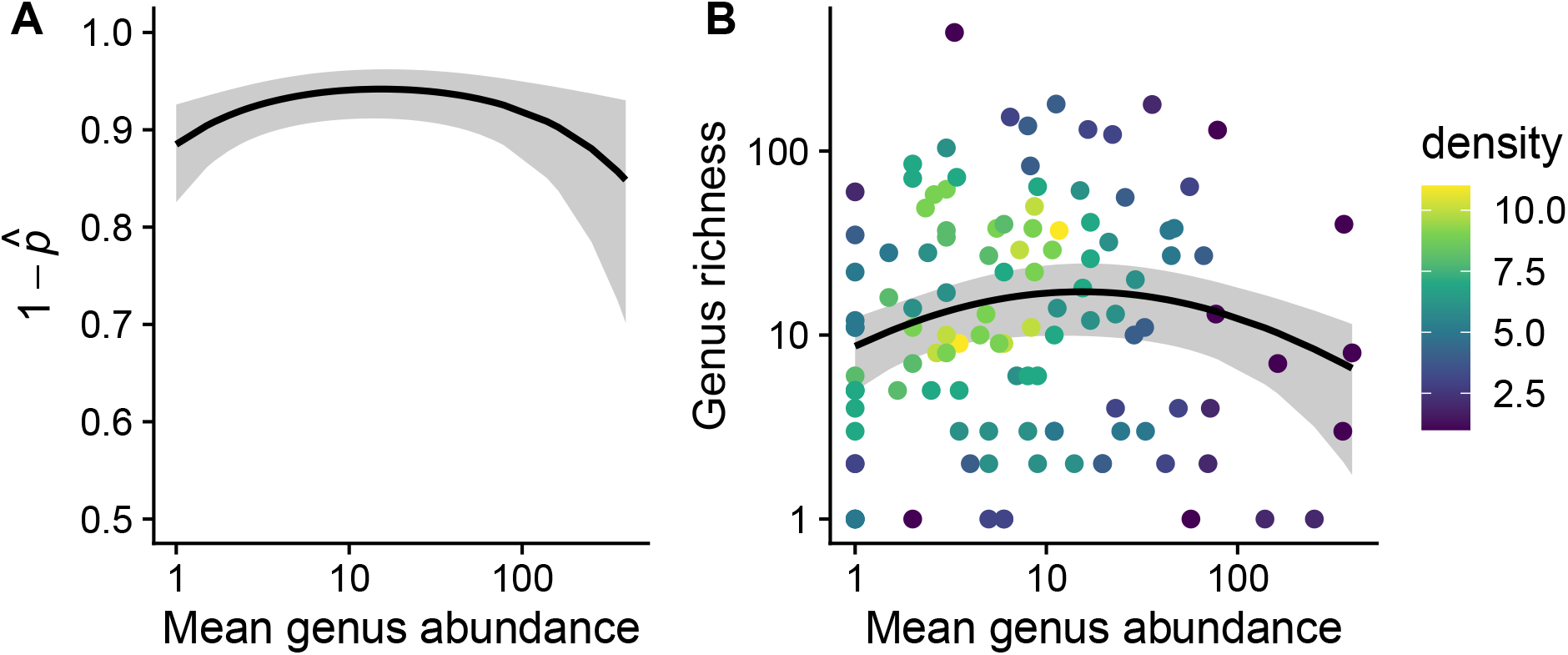
Relationship between genus abundance and species richness for arthropods endemic to Hawai’i. Panel A shows the fitted relationship between abundance and the observable diversification probability 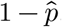. Solid line is expected value and gray shaded region is 95% confidence envelope. Panel B shows the pattern in the raw data along with the relationship predicted by the quadratic model. The solid black line represents the expected value and gray shaded regions the 95% confidence envelope. Colors of points reflect the local density of data points. The outlier with genus richness above 300 is *Drosophila* discussed further in the main text and Supplementary Section S5

The hump-shaped relationship is supported by a likelihood ratio test comparing quadratic and linear models of richness across abundance. The quadratic model is favored with a likelihood ratio of 7.012 and a *P*-value of 0.0081 from a χ^2^ distribution with *d*.*f*. = 1. Removing *Drosophila* has no meaningful impact on the relationship or the statistical significance of the quadratic term (see Supplementary Section S5).

## Discussion

Taken all together these results indicate support, from both theoretical and empirical perspectives, for the idea that intermediate abundances promote speciation. Using a theoretical simulation-based model we show that the chance of speciation peaks at intermediate abundances because these intermediate abundances represent a balance between dispersal, which slows speciation, and population persistence plus per capita incipient speciation, which are necessary for speciation. The balance arises because dispersal is lowest at low abundance (meaning the rate of transition from incipient to full speciation is greatest at low abundance) while persistence and incipient speciation are highest at high abundance. Our work suggests an intriguing possibility that if properties like abundance and dispersal are emergent species-level traits that modulate speciation, they could be acted upon by macroevolutionary selection (Rabosky and McCune 2010). Our results are novel in that they go against the widespread assumption that more abundant lineages should be those that produce more new speciation— an assumption tracing back to Darwin (1859) and also integral to the unified neutral theory of biodiversity (Hubbell 2001). However, our theoretical results are also grounded in published hypotheses, specifically the role of dispersal in modulating speciation (Price and Wagner 2004, Claramunt et al. 2012, Czekanski-Moir and Rundell 2019, Yamaguchi 2022).

Our empirical results support the theoretical finding that speciation peaks at intermediate abundances. The empirical results are made possible by the unique evolutionary setting of the Hawaiian Archipelago, where extreme isolation aids in the direct observation of diversification (Wagner and Funk 1995, Price and Wagner 2004). However, inferring past diversification from modern-only data is fraught (Gruner et al. 2008, Louca and Pennell 2020) and so we emphasize that the empirical results should be interpreted in concert with the theoretical results, each adding distinct and complimentary support to the idea of peak speciation at intermediate abundances.

The empirical and theoretical results are also distinct and cannot be fully interoperated. While the x-axis locations of the quadratic curve peaks in Figure 2 and Figure 3 are roughly similar, this does not mean that the simulation model parameters have a one-to-one mapping onto the parameters of a hypothetical diversification model that could have produced the empirical patterns. The parameters of the simulation model have to be interpreted separately from the empirical results because of key differences between the simulation set-up and the empirical analysis. Firstly, the “response variables” of each analysis are different—the simulation is able to directly observe speciation and thus we directly analyze speciation probability; conversely, we cannot directly observe speciation in the empirical system and so instead model 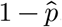, the “observable diversification probability.” The fact that we stop a simulation run once speciation is achieved also makes the simulated results different from the empirical system—in the real world, time does not stop if speciation occurs; instead, some clades continue to diversify and some go extinct. The continued diversification of real world clades could also explain the flatter relationship of 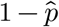 across abundances. Additionally, while both the simulation model and empirical analysis are concerned with abundance, how abundance is recorded in each is different. In the simulation we know the complete population size of every local population and we scale rates such that local population sizes are computationally tractable. Real world population sizes (perhaps up to 10^8^) would not be computationally manageable. The data we have also represent tiny subsamples of these populations.

Nonetheless, the simulation model provides a proof-of-concept and a mechanism for how speciation might depend on abundance in a more nuanced way than previously considered. Critically, our model does not depend on a neutrality assumption to arrive at this conclusion. Although our model is inspired by the agent-based modeling approaches of neutral theories (Hubbell 2001, Rosindell and Harmon 2013, Rosindell et al. 2015) we only model populations of a single species and thus make no assumption about how multiple species interact or whether different species have different eco-evolutionary rates. Thus our findings apply regardless of whether one assumes neutral- or niche-based dynamics. We also do not assume density dependence, but given that density-dependent dispersal (Travis et al. 1999) increases super-linearly with abundance, including density dependence would likely further accentuate our findings.

The mechanism agnostic approach of our model highlights the unnecessary assumptions of previous approaches. In particular Darwin (1859) assumed that abundant lineages were biologically dominant to less abundant lineages, and that this dominance promoted both abundance and diversification (Darwin 1859, Maurer 1999). A representative quote is reproduced below

Hence the struggle for the production of new and modified descendants will mainly lie between the larger groups, which are all trying to increase in number. One large group will slowly conquer another large group, reduce its numbers, and thus lessen its chance of further variation and improvement. Within the same large group, the later and more highly perfected sub-groups, from branching out and seizing on the many new places in the polity of Nature, will constantly tend to supplant and destroy the earlier and less improved sub-groups. Small and broken groups and sub-groups will finally tend to disappear. Looking into the future, we can predict that the groups which are now triumphant, and which as yet have suffered least extinction, will for a long period continue to increase. (Darwin 1859 pp. 125–26)

Darwin’s rhetoric about “conquer[ing]” and “triumphant” clades cannot be separated from discourse then and now that seeks to justify racism and colonial violence (Brantlinger 2003, Hesse 2007, Fuentes 2021, Diogo et al. 2023). Beyond the inherent amorality of such assumptions, our work shows them to be scientifically irrelevant, unnecessary, and incorrect in explaining how and why abundance connects to speciation.

## References

Afonso Silva, A. C., Maliet, O., Aristide, L., Nogués-Bravo, D., Upham, N., Jetz, W. and Morlon, H. 2025. Negative global-scale association between genetic diversity and speciation rates in mammals. - Nature communications 16: 1796.

Ashby, B., Shaw, A. K. and Kokko, H. 2020. An inordinate fondness for species with intermediate dispersal abilities. - Oikos 129: 311–319.

Birand, A., Vose, A. and Gavrilets, S. 2012. Patterns of species ranges, speciation, and extinction. - The American Naturalist 179: 1–21.

Brantlinger, P. 2003. Dark vanishings: Discourse on the extinction of primitive races, 1800–1930. - Cornell University Press.

Brown, J. H. 1995. Macroecology. - University of Chicago Press.

Chesson, P. 2000. Mechanisms of maintenance of species diversity. - Annual review of Ecology and Systematics 31: 343–366.

Ciccheto, J. R. M., Carnaval, A. C. and Araujo, S. B. L. 2024. The influence of fragmented landscapes on speciation. - Journal of Evolutionary Biology 37: 1499–1509.

Claramunt, S., Derryberry, E. P., Remsen Jr, J. and Brumfield, R. T. 2012. High dispersal ability inhibits speciation in a continental radiation of passerine birds. - Proceedings of the Royal Society B: Biological Sciences 279: 1567–1574.

Claramunt, S., Sheard, C., Brown, J. W., Cortés-Ramírez, G., Cracraft, J., Su, M. M., Weeks, B. C. and Tobias, J. A. 2025. A new time tree of birds reveals the interplay between dispersal, geographic range size, and diversification. - Current Biology 35: 3883–3895.

Cowie, R. H. and Holland, B. S. 2008. Molecular biogeography and diversification of the endemic terrestrial fauna of the hawaiian islands. - Philosophical Transactions of the Royal Society B: Biological Sciences 363: 3363–3376.

Czekanski-Moir, J. E. and Rundell, R. J. 2019. The ecology of nonecological speciation and nonadaptive radiations. - Trends in Ecology & Evolution 34: 400–415.

Darwin, C. 1859. On the origin of species by means of natural selection, or the preservation of favoured races in the struggle for life. - John Murray.

Diogo, R., Adesomo, A., Farmer, K. S., Kim, R. J. and Jackson, F. 2023. Not just in the past: Racist and sexist biases still permeate biology, anthropology, medicine, and education. - Evolutionary Anthropology: Issues, News, and Reviews 32: 67–82.

Eddelbuettel, D., Francois, R., Allaire, J., Ushey, K., Kou, Q., Russell, N., Ucar, I., Bates, D. and Chambers, J. 2025a. Rcpp: Seamless r and c++ integration.

Eddelbuettel, D., Francois, R., Bates, D., Ni, B. and Sanderson, C. 2025b. RcppArmadillo: 341 ‘Rcpp’ integration for the ‘Armadillo’ templated linear algebra library.

Etienne, R. S. and Haegeman, B. 2011. The neutral theory of biodiversity with random fission speciation. - Theoretical Ecology 4: 87–109.

Etienne, R. S., Apol, M. E. F., Olff, H. and Weissing, F. J. 2007. Modes of speciation and the neutral theory of biodiversity. - Oikos 116: 241–258.

Fuentes, A. 2021. “The descent of man,” 150 years on. - Science 372: 769–769.

Gaston, K. J. 2003. The structure and dynamics of geographic ranges. - Oxford University Press.

Gaston, K. J., Blackburn, T. M. and Lawton, J. H. 1997. Interspecific abundance-range size relationships: An appraisal of mechanisms. - Journal of animal Ecology: 579–601.

Goldberg, E. E., Lancaster, L. T. and Ree, R. H. 2011. Phylogenetic inference of reciprocal effects between geographic range evolution and diversification. - Systematic biology 60: 451–465.

Greenberg, D. A. and Mooers, A. Ø. 2017. Linking speciation to extinction: Diversification raises contemporary extinction risk in amphibians. - Evolution Letters 1: 40–48.

Gruner, D. S. 2007. Geological age, ecosystem development, and local resource constraints on arthropod community structure in the hawaiian islands. - Biological Journal of the Linnean Society 90: 551–570.

Gruner, D. S., Gotelli, N. J., Price, J. P. and Cowie, R. H. 2008. Does species richness drive speciation? A reassessment with the hawaiian biota. - Ecography 31: 279–285.

Hanski, I. 1998. Metapopulation dynamics. - Nature 396: 41–49.

Hay, E. M., McGee, M. D. and Chown, S. L. 2022. Geographic range size and speciation in honeyeaters. - BMC Ecology and Evolution 22: 86.

Hedges, S. B., Marin, J., Suleski, M., Paymer, M. and Kumar, S. 2015. Tree of life reveals clock-like speciation and diversification. - Molecular biology and evolution 32: 835–845.

Hesse, B. 2007. Racialized modernity: An analytics of white mythologies. - Ethnic and Racial Studies 30: 643–663.

Holt, R., Lawton, J., Gaston, K. and Blackburn, T. 1997. On the relationship between range size and local abundance: Back to basics. - Oikos: 183–190.

Hubbell, S. P. 2001. The unified neutral theory of biodiversity and biogeography. - Princeton University Press.

Irwin, J. O. 1975. The generalized waring distribution. Part i. - Journal of the Royal Statistical Society: Series A (General) 138: 18–31.

Jablonski, D. and Roy, K. 2003. Geographical range and speciation in fossil and living molluscs. - Proceedings of the Royal Society of London. Series B: Biological Sciences 270: 401–406.

Kendall, D. G. 1948. On the generalized” birth-and-death” process. - The annals of mathematical statistics: 1–15.

Krug, A. Z., Jablonski, D. and Valentine, J. W. 2008. Species–genus ratios reflect a global history of diversification and range expansion in marine bivalves. - Proceedings of the Royal Society B: Biological Sciences 275: 1117–1123.

Louca, S. and Pennell, M. W. 2020. Extant timetrees are consistent with a myriad of diversification histories. - Nature 580: 502–505.

Makarieva, A. M. and Gorshkov, V. G. 2004. On the dependence of speciation rates on species abundance and characteristic population size. - Journal of Biosciences 29: 119–128.

Matzke, N. J. 2014. Model selection in historical biogeography reveals that founder-event speciation is a crucial process in island clades. - Systematic biology 63: 951–970.

Maurer, B. A. 1999. Untangling ecological complexity: The macroscopic perspective. - University of Chicago Press.

Nee, S., May, R. M. and Harvey, P. H. 1994. The reconstructed evolutionary process. - Philosophical Transactions of the Royal Society of London. Series B: Biological Sciences 344: 305–311.

Nishida, G. M. 1992. Hawaiian terrestrial arthropod checklist. - Bishop University Press, Honolulu.

O’Grady, P. and DeSalle, R. 2018. Hawaiian drosophila as an evolutionary model clade: Days of future past. - Bioessays 40: 1700246.

Pennell, M. and MacPherson, A. 2025. Reading yule in light of the history and present of macroevolution. - Philosophical Transactions of the Royal Society B: Biological Sciences in press.

Price, J. P. and Wagner, W. L. 2004. Speciation in hawaiian angiosperm lineages: Cause, consequence, and mode. - Evolution 58: 2185–2200.

R Core Team 2025. R: A language and environment for statistical computing. - R Foundation for Statistical Computing.

Rabosky, D. L. and McCune, A. R. 2010. Reinventing species selection with molecular phylogenies. - Trends in Ecology & Evolution 25: 68–74.

Ricklefs, R. E. 2003. A comment on hubbell’s zero-sum ecological drift model. - Oikos 100: 185–192.

Rominger, A. J. 2026. abundolism.

Rosindell, J. and Harmon, L. J. 2013. A unified model of species immigration, extinction and abundance on islands. - Journal of Biogeography 40: 1107–1118.

Rosindell, J., Cornell, S. J., Hubbell, S. P. and Etienne, R. S. 2010. Protracted speciation revitalizes the neutral theory of biodiversity. - Ecology Letters 13: 716–727.

Rosindell, J., Harmon, L. J. and Etienne, R. S. 2015. Unifying ecology and macroevolution with individual-based theory. - Ecology letters 18: 472–482.

Smyčka, J., Toszogyova, A. and Storch, D. 2023. The relationship between geographic range size and rates of species diversification. - Nature Communications 14: 5559.

Stanley, S. M. 1986. Population size, extinction, and speciation: The fission effect in neogene bivalvia. - Paleobiology 12: 89–110.

Tao, R., Sack, L. and Rosindell, J. 2021. Biogeographic drivers of evolutionary radiations. - Frontiers in Ecology and Evolution 9: 644328.

Tilman, D. 1982. Resource competition and community structure. - Princeton university press.

Travis, J. M., Murrell, D. J. and Dytham, C. 1999. The evolution of density–dependent dispersal. - Proceedings of the Royal Society of London. Series B: Biological Sciences 266: 1837–1842.

Wagner, W. L. and Funk, V. A. 1995. Hawaiian biogeography: Evolution on a hot spot archipelago.

Yamaguchi, R. 2022. Intermediate dispersal hypothesis of species diversity: New insights. - Ecological Research 37: 301–315.

Yee, T. W. 2025. VGAM: Vector generalized linear and additive models.

Yule, G. U. 1925. A mathematical theory of evolution, based on the conclusions of dr. JC willis, FR s. - Philosophical transactions of the Royal Society of London. Series B, containing papers of a biological character 213: 21–87.

