## Supplement for "Intermediate abundance promotes speciation when dispersal is limited"

### S1 Computation set-up

We make use of a custom R package *abundolism* (Rominger 2026) which can be installed from github.

```
devtools::install_github("ajrominger/abundolism")
```

The following R computing environment is used for all simulations and analyses:

```
library(abundolism)
library(dplyr)
```

```
Attaching package: 'dplyr'

The following objects are masked from 'package:stats':

    filter, lag

The following objects are masked from 'package:base':

    intersect, setdiff, setequal, union
```

```r
library(ggplot2)
library(ggpointdensity)
library(cowplot)
library(knitr)

sessionInfo() |>
    print(locale = FALSE)
```

```
R version 4.5.2 (2025-10-31)
Platform: aarch64-apple-darwin20
Running under: macOS Sequoia 15.0.1

Matrix products: default
BLAS:   /System/Library/Frameworks/Accelerate.framework/Versions/A/Frameworks/vecLib.framewo[illegible]
LAPACK: /Library/Frameworks/R.framework/Versions/4.5-arm64/Resources/lib/libRlapack.dylib;  [illegible]

attached base packages:
[1] stats     graphics  grDevices utils     datasets  methods   base

other attached packages:
[1] knitr_1.50          cowplot_1.2.0.9000   ggpointdensity_0.2.1
[4] ggplot2_4.0.1       dplyr_1.2.1          abundolism_0.1.0

loaded via a namespace (and not attached):
 [1] vctrs_0.7.3        cli_3.6.5          rlang_1.2.0        xfun_0.52
 [5] generics_0.1.4     S7_0.2.1           jsonlite_2.0.0     glue_1.8.0
 [9] htmltools_0.5.8.1  scales_1.4.0       rmarkdown_2.29     grid_4.5.2
[13] evaluate_1.0.4     tibble_3.3.0       MASS_7.3-65        fastmap_1.2.0
[17] yaml_2.3.10        lifecycle_1.0.5    compiler_4.5.2     RColorBrewer_1.1-3
[21] Rcpp_1.1.0         pkgconfig_2.0.3    rstudioapi_0.17.1  farver_2.1.2
[25] digest_0.6.39      R6_2.6.1           tidyselect_1.2.1   pillar_1.11.0
[29] magrittr_2.0.4     withr_3.0.2        tools_4.5.2        gtable_0.3.6
```

The quarto (Allaire et al. 2025) document used to generate this supplement can be found with the R command `system.file("ab_supp.qmd", package = "abundolism")`.

### S2 Additional simulation details

The birth-death-immigration-speciation simulation tracks the following:

- all local population sizes
- running mean population size per iteration
- time relative to all rates (i.e. not simply number of iterations)
- whether an incipient speciation event has occurred
- the wait time to full speciation
- whether full speciation has occurred

In each iteration of the simulation an event is chosen at random. Possible events for each local population are

- local birth
- local death
- local speciation
- local immigration

There is also one possible global event: immigration from the source pool.

If there are $n_p$ local populations then the total number of possible events are $n_p \times 4 + 1$. Which event occurs during any given simulation iteration is chosen at random according to the probability of each event. With a local population size of $x_i$, the probability of each event is proportional to its respective rate:

$$\mathbb{P}(\text{birth in pop. } i) = \frac{\lambda x_i}{R} \tag{1}$$

$$\mathbb{P}(\text{death in pop. } i) = \frac{\mu x_i}{R} \tag{2}$$

$$\mathbb{P}(\text{speciation in pop. } i) = \frac{\nu x_i}{R} \tag{3}$$

$$\mathbb{P}(\text{local immigration from pop. } i) = \frac{m_{prop} \gamma x_i}{R} \tag{4}$$

$$\mathbb{P}(\text{global immigration}) = \frac{\gamma}{R} \tag{5}$$

where $R = \gamma + \left(\sum_i^{n_p} x_i\right)(\lambda + \mu + \nu + m_{prop}\gamma)$ is a normalization constant.

In the case of immigration (both local and from the source pool) the recipient of the dispersing individual must also be chosen. We choose the recipient local population at random with equal probability across local populations.

Once an event is randomly sampled, the variables being tracked are updated. Birth and immigration (local or global) augment a local population by 1. Death decrements a local population by 1. Immigration events have no impact on the source of the dispersing individual. Time is updated following the Gillespie algorithm: the sojourn time between events is a random variable with an exponential probability distribution with rate R: $s \sim Exp(R)$. Each time step a random sojourn time is sampled and added to the running total time since the start of the simulation.

If a population experiences local extirpation all of its speciation-related information is re-set including removing any incipient speciation and restoring the wait time to full speciation back to the default of $\tau$.

If full speciation is achieved the simulation stops. If no speciation occurs but the specified number of simulation iterations is reached, the simulation stops. Once stopped, the simulation returns the tracked variables of time, mean population size, and whether full speciation has occurred.

### S3 Simulation experiment details

Simulation parameters are drawn from uniform distributions with the following minimum and maximum limits:

```r
par_range <- data.frame(
    param = c("la", "mu", "g", "m_prop", "nu", "tau", "xi"),
    min   = c(0.1,  0.1,  0.1,  0,        0,     20,  10),
    max   = c(4,    4,    1,    0.1,      0.1,  200, 100)
)


# fixed params
np <- 10
nstep <- 10000

# number of simulation replicates
nrep <- 5000

# format param names for printing
mutate(par_range,
       param = sprintf("`%s`", param)) |>
    kable(format.args = list(scientific = FALSE))
```

| param | min | max |
|---|---|---|
| `la` | 0.1 | 4.0 |
| `mu` | 0.1 | 4.0 |
| `g` | 0.1 | 1.0 |
| `m_prop` | 0.0 | 0.1 |

| param | min | max |
|---|---|---|
| nu | 0.0 | 0.1 |
| tau | 20.0 | 200.0 |
| xi | 10.0 | 100.0 |

The parameters governing number of populations (`np`) and number of time steps (`nstep`) do not vary across simulations and are set to `np = 10` and `nstep = 10`$^4$. We simulate a total of `nrep = 5000` replicates, each with a unique, randomly sampled set of parameter values.

#### S3.1 Running the simulation experiment

Parameter values for each simulation replicate are stored in a `data.frame` with one column for each parameter.

```
# change or remove for different results
set.seed(123)

# data.frame of randomly sampled param values
# one param per column
pars <- sapply(par_range$param, function(p) {
    runif(nrep,
          par_range[par_range$param == p, "min"],
          par_range[par_range$param == p, "max"])
}) |>
    as.data.frame()
```

With those parameters sampled, we can now run the simulation, one realization for each parameter combination. For details on inputs and outputs of the simulation function `sim_BDI_spec` see the help documentation accessible with `?sim_BDI_spec`.

```
# now run the simulation
sim_dat <- sim_BDI_spec(la = pars$la, mu = pars$mu, g = pars$g,
                        m_prop = pars$m_prop, nu = pars$nu,
                        tau = pars$tau, xi = pars$xi,
                        np = np, nstep = nstep)
```

### S4 Interpreting the simulation experiment

Now we can visually inspect the results. Our main purpose is to study how abundance relates to speciation in this model, but first we check other relationships to ensure the model behaves as expected. In particular, population size should be larger when $\lambda > \mu$ and smaller, supported primarily by immigration. When $\lambda < \mu$ populations will grow indefinitely. When birth and death rates are equal ($\lambda = \mu$) the population could crash or expand depending on small random fluctuations and random immigration. Supplementary Figure S1 shows exactly this pattern. Of note: simulation runs experiencing speciation diverge from the expected pattern but that is specifically because our simulation algorithm stops the simulation short (i.e. populations are not allowed to continue changing) once speciation occurs.

```
ggplot(sim_dat, aes(la - mu, mean_pop_size)) +
    geom_point(data = select(sim_dat, !speciation),
               mapping = aes(la - mu, mean_pop_size), color = "gray50") +
    geom_pointdensity() +
    facet_wrap(vars(speciation),
               labeller = as_labeller(c(`0` = "No speciation",
                                        `1` = "Yes speciation"))) +
    scale_y_log10() +
    xlab(expression(italic(lambda - mu))) +
    ylab("Mean population size") +
    scale_color_viridis_c(trans = "log10") +
    theme_cowplot()
```

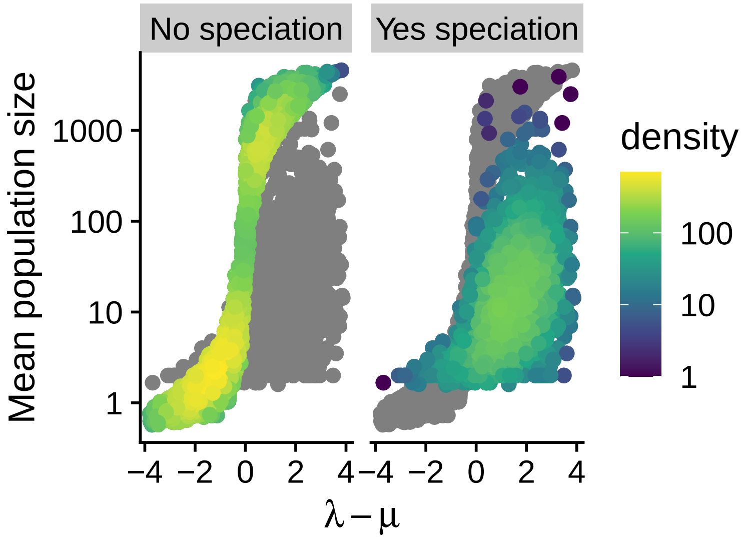


Supplementary Figure S1: Relationship across simulation runs between growth rate ($\lambda - \mu$) and mean population size. Panels show simulation runs without and with speciation. Points are color coded according to local density of points in an attempt to more clearly show the distribution of simulation results.

```r
# model of speciation changing with nu
nu_mod <- glm(speciation ~ nu, data = sim_dat, family = binomial)
nu_slope_ci <- confint(nu_mod, 2)
```

We also expect that probability of specification should increase with the parameter $\nu$, the rate of incipient speciation. Supplementary Figure S2 confirms this expectation but shows the relationship to be weak, indicating that while this parameter obviously matters, its effect is modulated by other factors such as abundance and dispersal. A logistic regression model confirms this weakly positive but meaningful relationship estimating a 95% confidence interval on the slope of (1.512, 5.748).

With results from the simulation we report confidence intervals rather than hypothesis tests because sample sizes from simulations are arbitrary and thus degrees of freedom can be manipulated to produce statistically significant results from biologically meaningless patterns. By contrast, confidence intervals more meaningfully indicate the magnitude of effects in the simulation experiment.

```r
spec_plot(sim_dat, nu,
          xlab = expression("Incipient speciation rate"~italic(nu)))
```

```
`geom_smooth()` using formula = 'y ~ x'
```

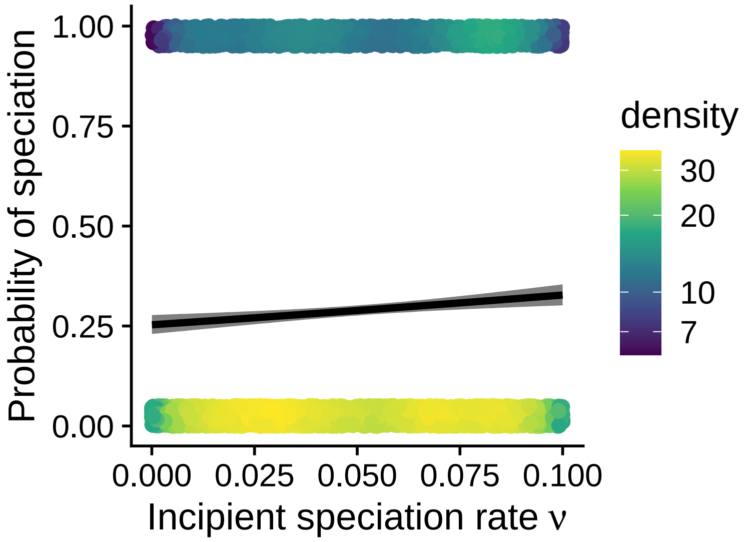


Supplementary Figure S2: Relationship across simulation runs between incipient speciation rate and speciation. Points are jittered around 1 and 0 and are color coded according to local density of points all in an attempt to more clearly show the distribution of simulation results. Curve is a binomial generalize linear model and gray region shows 95% confidence envelope.

```
# model of speciation changing with dispersal
disp_mod <- glm(speciation ~ I(log((g + m_prop * g) / la)),
                data = sim_dat, family = binomial)
disp_slope_ci <- confint(disp_mod)
```

We also expect dispersal to decrease the chance of speciation because dispersal prolongs the wait time until full speciation. But we have to evaluate this expectation with respect to *relative* dispersal, that is dispersal relative to birth because these rate trade-off in determining whether a population grows due to intrinsic (birth) or extrinsic (dispersal) forces. Dispersal comes from two sources: global dispersal and local dispersal, so we add these two sources when considering total dispersal. Supplementary Figure S3 Confirms the negative relationship between speciation and dispersal. The slope from a logistic regression model for log dispersal normalized by birth rate has a 95% confidence interval of (-1.695, -0.602).

```
spec_plot(sim_dat, (1 + m_prop) * g / la, xlog = TRUE,
          xlab = "Relative dpeciation rate")
```

```
`geom_smooth()` using formula = 'y ~ x'
```

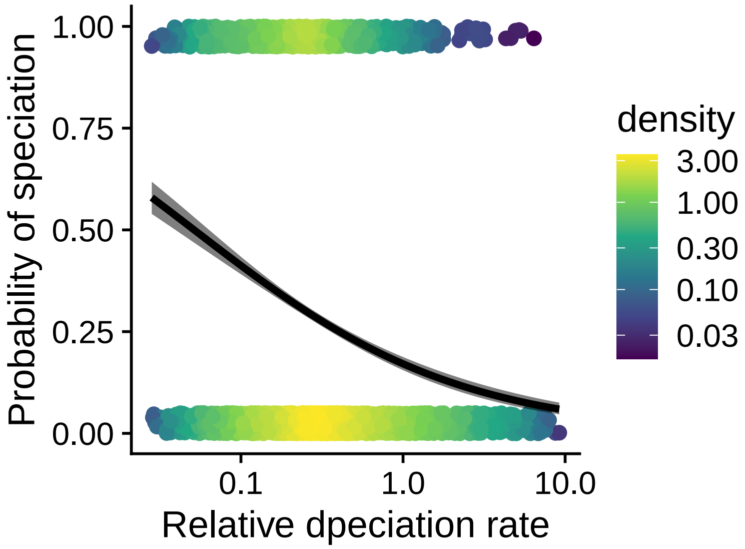


Supplementary Figure S3: Relationship across simulation runs between relative dispersal rate and speciation. Points are jittered around 1 and 0 and are color coded according to local density of points all in an attempt to more clearly show the distribution of simulation results. Curve is a binomial generalize linear model and gray region shows 95% confidence envelope.

```
# model of speciation changing with wait time to full speciation
tau_mod <- glm(speciation ~ log(tau),
               data = sim_dat, family = binomial)
tau_slope_ci <- confint(tau_mod, 2)
```

We might also expect that speciation decreases with the duration of the wait time $\tau$ to full speciation. However, $\tau$ is only the initial value for the wait time, dispersal continually augments this wait time, thus $\tau$ by itself does not impact probability of speciation. The 95% confidence interval for the slope of speciation probability against $\log(\tau)$ is (-0.132, 0.073), which does skew toward negative values, but includes the value of slope = 0.

#### S4.1 Evaluating the relationship between abundance and speciation

These results are reported in the main text. Here we show how the results from the main text are derived.

First we build a quadratic model of probability of speciation as a function of abundance with the below code:

```
sim_dat$log_abund <- log(sim_dat$mean_pop_size)
sim_mod <- glm(speciation ~ log_abund + I(log_abund^2),
               family = binomial, data = sim_dat)

sim_quad_ci <- confint(sim_mod, parm = 3)
```

```
Waiting for profiling to be done...
```

The confidence interval for the quadratic term is far from 0:

```
sim_quad_ci
```

```
     2.5 %     97.5 %
-0.4156320 -0.3651633
```

We now turn to developing the figure of abundance versus speciation. These results are reported in the main text, but the code below is used to create the visualization of speciation versus abundance.

```
special_log10_breaks <- function(x) {
    p <- seq(floor(x[1]),
             ceiling(x[2]),
             by = (floor(x[2]) - ceiling(x[1])) / 15)

    10^p
}

spec_by_bin <- mutate(sim_dat,
                      bins = cut(
                          mean_pop_size,
                          log(mean_pop_size, 10) |>
                              range() |>
                              special_log10_breaks()
                      )) |>
    summarize(spec_prop = mean(speciation),
              n = n(),
              .by = bins) |>
    mutate(xlo = gsub("\\(|,.*", "", bins) |> as.numeric(),
           xhi = gsub("\\]|.*,", "", bins) |> as.numeric()) |>
    arrange(xlo)
```

```r
# plotting
sim_fig <- spec_plot(sim_dat, mean_pop_size, xlab = "Mean population size",
               xlog = TRUE, formula = y ~ x + I(x^2),
               se = FALSE) +
    geom_rect(data = spec_by_bin,
              mapping = aes(xmin = xlo, xmax = xhi,
                            ymin = 0, ymax = spec_prop),
              alpha = 0.25,
              inherit.aes = FALSE)

# change order of layer display
sim_fig$layers <- sim_fig$layers[c(1, 3, 2)]

# add background bars
sim_fig <- sim_fig +
    theme(panel.background = element_rect(fill = "gray90"),
          panel.grid.major.x = element_line(color = "white"),
          panel.grid.minor.x = element_line(color = "white")) +
    scale_x_log10(minor_breaks = spec_by_bin$xlo, expand = FALSE)
```

```
Scale for x is already present.
Adding another scale for x, which will replace the existing scale.
```

### S5 Analyzing abundance and richness of arthropod genera in Hawai‘i

The results of analyzing Hawai‘i arthropod data are presented in the main text. The code to produce these results is shown here. First we load and prepare the data. Preparation involves filtering data to only endemic taxa and compiling mean abundance for genera based on data from Gruner (2007) and compiling species richness for general based on the Bernice Pauahi Bishop Museum arthropod checklist (Nishida 1992).

```r
# read gruner data
arth <- read.csv("raw_data/arth.csv")

# read checklist data
arth_check <- read.csv("raw_data/hawaii_arthropod_checklist.csv")

# calculate richness of genera from checklist
```

```
gen_rich <- filter(arth_check, Genus != "") |>
    group_by(Genus) |>
    summarize(nspp = n_distinct(Genus.and.species)) |>
    ungroup() |>
    rename(genus = Genus)

# calculate abundances of genera from survey data
gen_abund <- filter(arth, origin == "native-endemic") |>
    group_by(genus, species_code) |>
    summarize(abund = n()) |>
    ungroup(species_code) |>
    summarize(mean_abund = mean(abund)) |>
    ungroup()
```

```
`summarise()` has grouped output by 'genus'. You can override using the
`.groups` argument.
```

```
# `gen` holds data ready to analyze for richness ~ abundance
gen <- inner_join(gen_rich, gen_abund)
```

```
Joining with `by = join_by(genus)`
```

With the prepared data we can compute a quadratic and linear model of genus richness versus abundance. As explained in the main text, these models are beta-binomial generalized linear models and are achieved with custom functions available in the *abundolism* package (Rominger 2026) associated with this paper. The main function is `glm_bgeom` which maximizes the log likelihood function and shares some common methods with `lm` and `glm` including `logLik` which enables likelihood analyses such as the likelihood ratio test.

To summarize from the main text, our full quadratic model is

$$\begin{aligned} y_i &\sim \text{BetaGeometric}(\alpha_i, \beta) \\ \log(\alpha_i) &= a_0 + a_1 x_i + a_2 x_i^2 \\ \log(\beta) &= b_0 \end{aligned}$$

where $x_i$ is mean abundance of lineage $i$ and our estimate for the observable diversification probability for lineage $i$ is $1 - \hat{p}_i = 1 - \frac{\alpha_i}{\alpha_i + \beta}$.

```
# add column for log abundance
gen$log_abund <- log(gen$mean_abund)

# quadratic model
arth_mod <- glm_bgeom(nspp ~ log_abund + I(log_abund^2), data = gen)

# straight line model
arth_mod0 <- glm_bgeom(nspp ~ log_abund, data = gen)

# view the model output
arth_mod
```

```
Call:  glm_bgeom(formula = nspp ~ log_abund + I(log_abund^2), data = gen)

Coefficients:
     (Intercept)        log_abund   I(log_abund^2) (Intercept):beta
       0.9204501       -0.5453639        0.1000263        2.9611979

Log likelihood:
'log Lik.' -493.7613 (df=4)
```

```
arth_mod0
```

```
Call:  glm_bgeom(formula = nspp ~ log_abund, data = gen)

Coefficients:
     (Intercept)        log_abund (Intercept):beta
       0.4126521       -0.0545682        2.7655422

Log likelihood:
'log Lik.' -497.2671 (df=3)
```

The likelihood ratio test comparing quadratic and linear models is achieved with a custom function `lrt_bgeom` from the *abundolism* package.

```
arth_lrt <- lrt_bgeom(arth_mod0, arth_mod)
```

To produce visualizations of the arthropod data we make use of a custom `ci_bgeom` function from *abundolism* that calculates confidence intervals and predictions from the output of `glm_bgeom`. This function first calculates standard errors for the $\log(\alpha)$ and $\log(\beta)$ terms of the beta-geometric GLM using the Wald approximation and then computes the standard error for $1 - \hat{p}$ using a change of variable approach. The predicted value $\hat{y}$ for numbers of species per genus is then $\hat{y} = \frac{1}{1-\hat{p}}$ and its standard error is calculated from that of $1 - \hat{p}$ again with a change of variable approach.

```
arth_mod_ci <- cbind(mean_abund = gen$mean_abund,
                     ci_bgeom(arth_mod, type = "response"))

arth_p_ci <- cbind(mean_abund = gen$mean_abund,
                   ci_bgeom(arth_mod, type = "prob"))

gp_nspp <- ggplot(arth_mod_ci, aes(mean_abund, ymin = y_lwr, ymax = y_upr)) +
    geom_ribbon(fill = "gray60", alpha = 0.5) +
    scale_x_log10() +
    scale_y_log10() +
    xlab("Mean genus abundance") +
    ylab("Genus richness") +
    geom_pointdensity(data = gen, mapping = aes(mean_abund, nspp),
                      inherit.aes = FALSE) +
    scale_color_viridis_c() +
    geom_line(data = arth_mod_ci,
              mapping = aes(mean_abund, y_hat),
              linewidth = 1) +
    theme_cowplot()


gp_prb <- ggplot(arth_p_ci, aes(mean_abund, y = p_hat,
                                ymin = p_lwr, ymax = p_upr)) +
    geom_ribbon(fill = "gray60", alpha = 0.5) +
    geom_line(linewidth = 1) +
    scale_x_log10() +
    ylim(0.5, 1) +
    xlab("Mean genus abundance") +
    ylab(expression(italic(1 - hat(p)))) +
    theme_cowplot()

gp_arth <- plot_grid(gp_prb, gp_nspp, rel_widths = c(0.75, 1),
                     labels = c("A", "B"), label_x = 0.02)
```

#### S5.1 Evaluating impact of outliers

There is one clear outlier: *Drosophila* with high richness given abundance. To make sure this outlier does not bias results we re-run the analysis of genus richness versus abundance without *Drosophila*.

```
arth_nodro <- glm_bgeom(nspp ~ log_abund + I(log_abund^2),
                        data = gen[gen$nspp < 400, ])
arth_nodro0 <- glm_bgeom(nspp ~ log_abund,
                         data = gen[gen$nspp < 400, ])
```

Based on a likelihood ratio test we find the quadratic model is still preferred.

```
lrt_bgeom(arth_nodro0, arth_nodro)
```

```
Likelihood ratio test

Model 0: nspp ~ log_abund
Model 1: nspp ~ log_abund + I(log_abund^2)

        #Df  LogLik Df  Chisq Pr(>Chisq)
Model 0   3 -486.59
Model 1   4 -483.10  1 6.9828    0.00823 **
---
Signif. codes:  0 '***' 0.001 '**' 0.01 '*' 0.05 '.' 0.1 ' ' 1
```

### References

Allaire, J. J., Teague, C., Scheidegger, C., Xie, Y., Dervieux, C. and Woodhull, G. 2025. Quarto.

Gruner, D. S. 2007. Geological age, ecosystem development, and local resource constraints on arthropod community structure in the hawaiian islands. - Biological Journal of the Linnean Society 90: 551–570.

Nishida, G. M. 1992. Hawaiian terrestrial arthropod checklist. - Bishop University Press, Honolulu.

Rominger, A. J. 2026. abundolism.
